# Importance of Matrix-dimensionality in Regulating the Bone Marrow Hematopoietic Cells Pool

**DOI:** 10.1101/720359

**Authors:** P. Zhang, C. Zhang, J. Han, J. Gao, W. Zhao, H. Yang

## Abstract

In bone marrow, hematopoietic stem cells (HSCs) and multiple hematopoietic progenitor cells (HPCs) cooperate to differentiate and replenish blood and immune cells. It has long been recognized bone marrow niche parameters interact with hematopoietic stem and progenitor cells (HSPCs) and additional work is required to study niche physical signals controlling cell behavior. Here we presented that important biophysical signals, stiffness and dimensionality, regulating expansion of bone marrow HSPCs. Mice bone marrow derived progenitor cells were cultured in collagen I hydrogel in vitro. We found stiffer 3D matrix promoted the expansion of lineage negative (Lin^−^) progenitor cells and Lin^−^Sca-1^+^c-kit^+^ (LSK) HSPCs compared to softer hydrogel. Compared with cells cultured in 2D environment, 3D embedded construct had significant advantage on HSPCs expansion, accompanied by increases on myeloid and lymphoid lineage fractions. Bright changes on gene expression were subsequently discovered. According to these data, we concluded that culture matrix dimensionality is an important factor to regulate the behavior of subpopulations in hematopoietic cell pool, which should be considered in attempts to illuminate HSCs fate decision in vitro.

**Statement of Significance:** We would like to submit the enclosed manuscript entitled "Importance of Niche-dimensionality in Regulating the Bone Marrow Hematopoietic Cells Pool", which we wish to be considered for publication in Biophysical Journal. Studies about the interaction between HSCs and factors provided by their microenvironment is largely focus on pure perspective of biology. But biophysical factors affecting HSC fate and behavior needs to be further explore. Herein we found ex vivo culture dimensionality affected HSPC expansion. Cell surface marker detection and mRNA expression analysis predicted the changes on myeloid and lymphoid lineage fractions. We hope niche physical signals which we identified will be considered to design HSC biomimetic niches in clinical applications. And we believe that our study will make it interesting to general readers. We deeply appreciate your consideration of our manuscript, and we look forward to receiving comments from the reviewers.

---

Research in stem cell biology relies on the knowledge of the cell microenvironment in vivo, known as “stem cell niche”, where stem cells are nurtured by the niche signals. A diverse array of niche parameters contributing to stem cell fate afford references for ex vivo niche mimicking and tissue engineering(1). So far we have some understanding of the interaction between cells and their niche components, such as stromal cells, soluble factors they produce, and matrix (ECM) molecules. But because of complexity and tissue specificity of niche, other requisite interacting factors need to be explored and require further characterization in each tissue(2). Hematopoietic stem cells (HSCs) locate at the top of hematopoietic hierarchy, is progenitor of all mature blood cells. HSC transplantation (HSCT) remains until now the only proven clinical application of stem cells and offers hope on hematologic disease treatment(3–5). In adult, a pool of hematopoietic cells, including HSCs, primarily reside in the bone marrow (BM) niches. HSCs receive important signals from niche microenvironment to maintain cell behaviors including quiescence, self-renew, proliferation and differentiation. Therefore, thoroughly understanding complexity of the niche facilitates engineering on artificial models to enable HSC research in vitro(6). Associations among HSCs biology and cellular constituents or biomolecular within BM cavity have been presented in quantity(7). Recently, evidence that spatiotemporal biophysical characteristics of the specific niche regulatory mechanism are increasing revealed, such as ECM matrix stiffness, dimensionality, and geometry/topography properties(8, 9). Herein we constructed collagen I hydrogel culture systems and presented that matrix stiffness and dimensionality are important biophysical signals regulating expansion of hematopoietic stem and progenitor cells (HSPCs) in BM hematopoietic cell pool. Perceive of BM niche biophysical characteristics is expected to accelerate novel remarkable advance regarding engineered cell culture environment.

Elastic moduli (stiffness) of hydrogel matrix with different collagen I concentration exhibited significantly difference (Figure 1A). Freshly isolated cells were cultivated within 3D collagen gel. Flow cytometric analysis revealed 3D substrates stiffness affected expansion of hematopoietic cells. Stiffer hydrogel promoted expansion of Lin^−^ progenitor cells and LSK HSPCs. Percentage of Lin^−^ cells within stiffer matrix was significantly raised from 22.00% to 25.93%. Stiffer culture system increased the percentage of LSK cells from 1.15% to 1.98%. Although expansion of LSK cells was clearly elevated, the statistic didn’t exhibit significant difference (Figure 1B).

**Figure 1.**
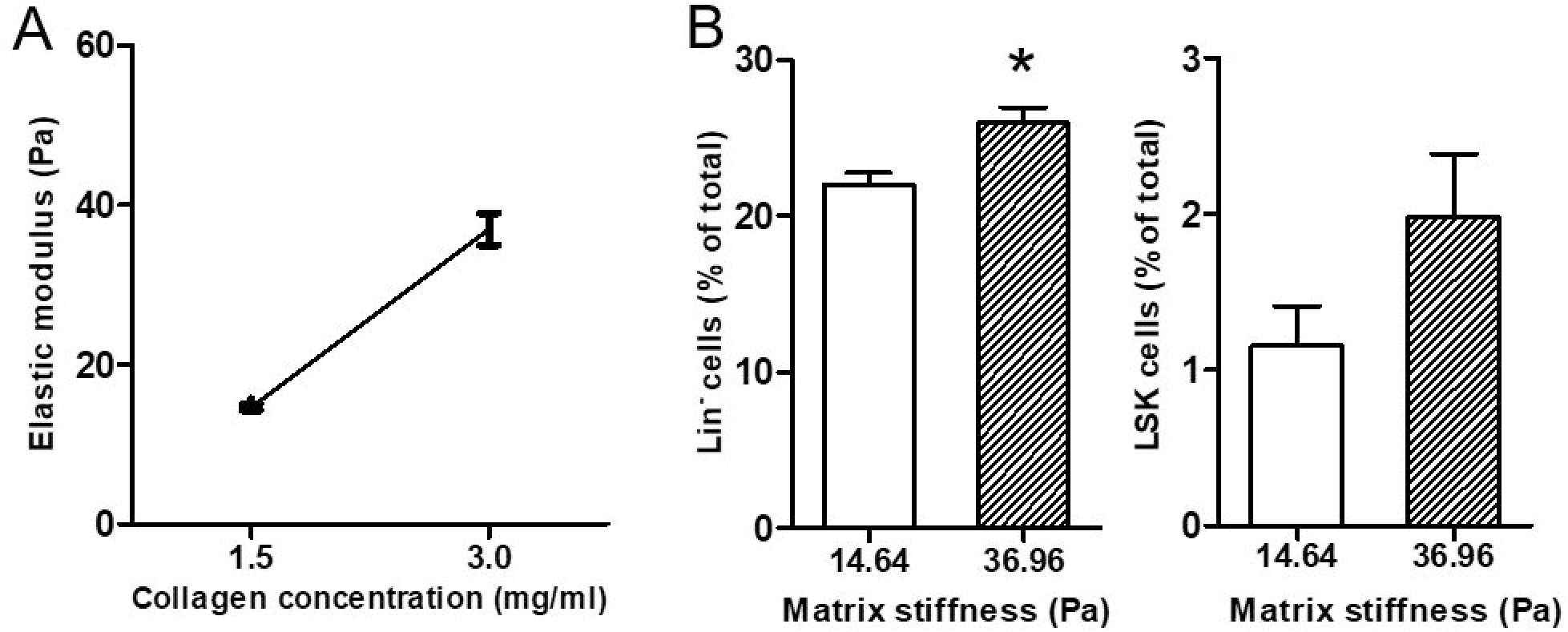
Stiffness of 3D hydrogel system affects expansion of mice HSPCs. (A) Stiffness of the hydrogel substrates used in the experiments. The substrate elastic moduli showed 14.65 Pa and 36.96Pa. n=5~7. Mean ± SEM. (B) Effects of culture system stiffness on expansion of Lin^−^ cells and LSK cells. Significant differences were observed on change of Lin^−^ fraction (p=0346). n=3. Mean ± SEM. *p<0.05.

To evaluate the impact of matrix-dimensionality on hematopoietic cells, cells were harvested from top of hydrogel matrix and 3D embedded construct fabricated by 3mg/ml collagen I for subsequent analysis (Figure S1). We discovered expansion of Lin^−^ cells as well as LSK cells were both enforced significantly by 3D matrix, in relative to the percentages of those on gel surface (Figure 2, 2S). Proportion of Lin^−^ cells in 2D environment was 5.72% and LSK cells was 0.24%. Within 3D system, proportion of Lin^−^ cells was distinctly improved to 15.00%. LSK frequency was increased to 1.033% (Figure2 B, C). To explain whether matrix-dimensionality influence the survival of myeloid and lymphoid lineage cells, we detected frequency changes on CD11b^+^ cells and CD19^+^ cells, representing myeloid and lymphoid cells in BM cell pool, respectively(Figure 3). Hematopoietic cells that were 3D embedded had increased myeloid (57.63% to 62.43%) lineage (Figure 3A). Lymphoid cells increased from 16.07% to 22.27% (Figure 3B). Both advances on myeloid and lymphoid fraction were observed with statistical significance. mRNA level analysis were performed to measure gene factors that participate in hematopoiesis development and hematopoietic cells function, containing HSC fingerprints *Gata2* and *Fgd5*, HSC homing related genes *Cxcr4* and *Cxcl12*, HSC differentiated fingerprint *Cd48*, lymphoid fingerprints *Atm* and *Fgf13*, myeloid fingerprints *Tlr13* and *Trem1*, and niche signals related genes *Wnt3a* and *Notch1*(10). Gene expression alternations suggested changes on HSPC behaviors including differentiation and mobilization led by matrix-dimensionality (Figure 3C). Elative gene expression levels of Gata2 and Fgd5 required for HSC generation and survival were overexpressed in 3D matrix. Specifically, we found 3D environment severely enhanced the fold change of *Trem1*, which is associated with myeloid lineage development. Up-regulation of gene expression of Wnt3a and Notch1 represented that canonical Wnt and Notch signaling in hematopoietic cells could be motived by the matrix dimensionality. Downward trend of Cxcl12 predicted 3D system possibly attenuated the cell migration.

**Figure 2.**
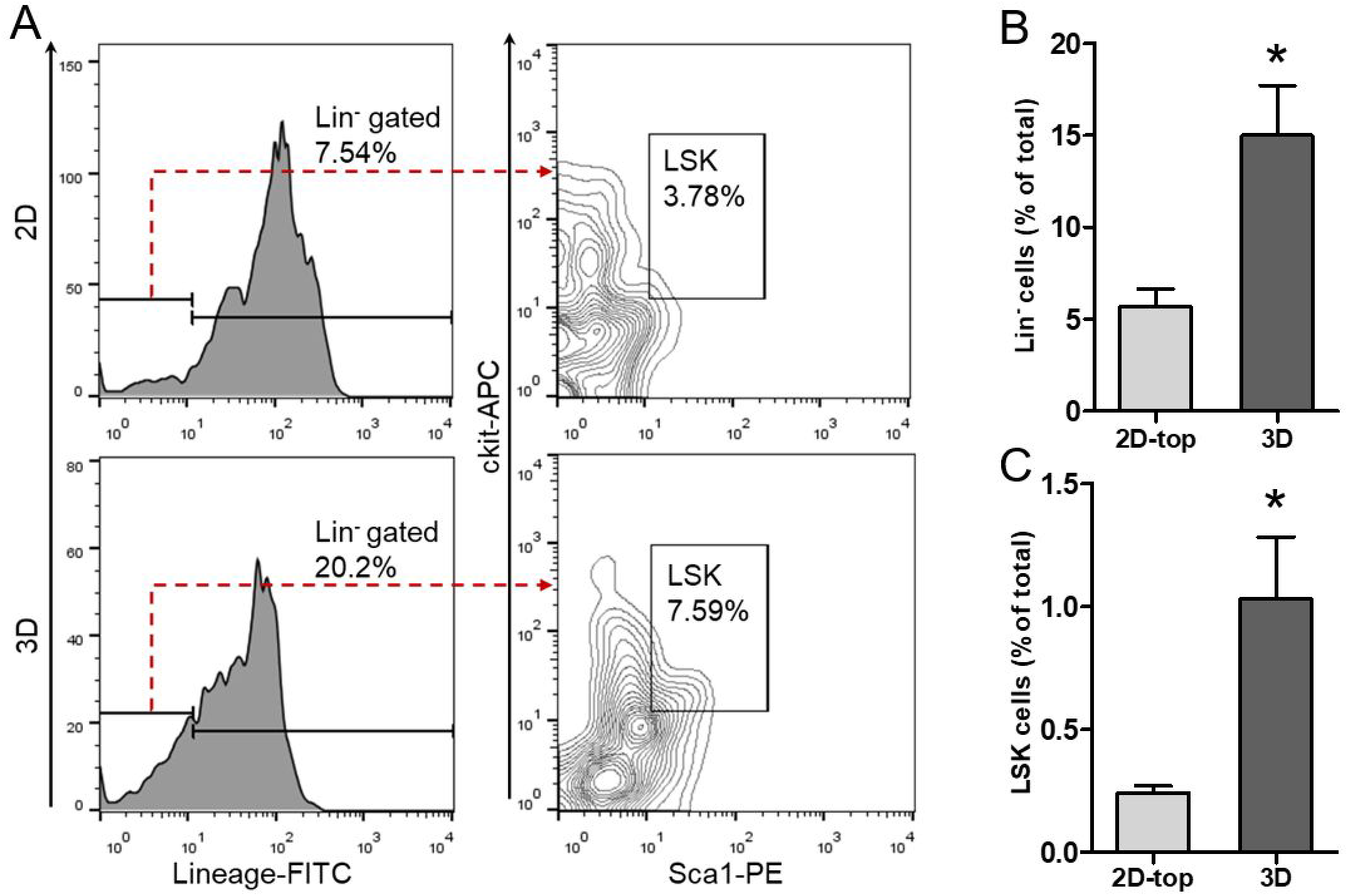
Effect comparison of matrix dimensionality (2D versus 3D) on expansion of HSPCs. (A) Flow cytometry assay showed the fraction of Lin^−^ progenitor cells (FITC negative) and LSK HSPCs (PE positive and APC positive population in Lin^−^ gated cells). (B) Lin^−^ cells and LSK cells percentages of both culture systems were compared according to flow cytometry analysis. 3D environment promoted extension of HSPCs rather than 2D. Differences were statistically significant (Lin^−^:p=0.0314, LSK:p=0.0351). n=5. Mean ± SEM. *p<0.05.

**Figure 3.**
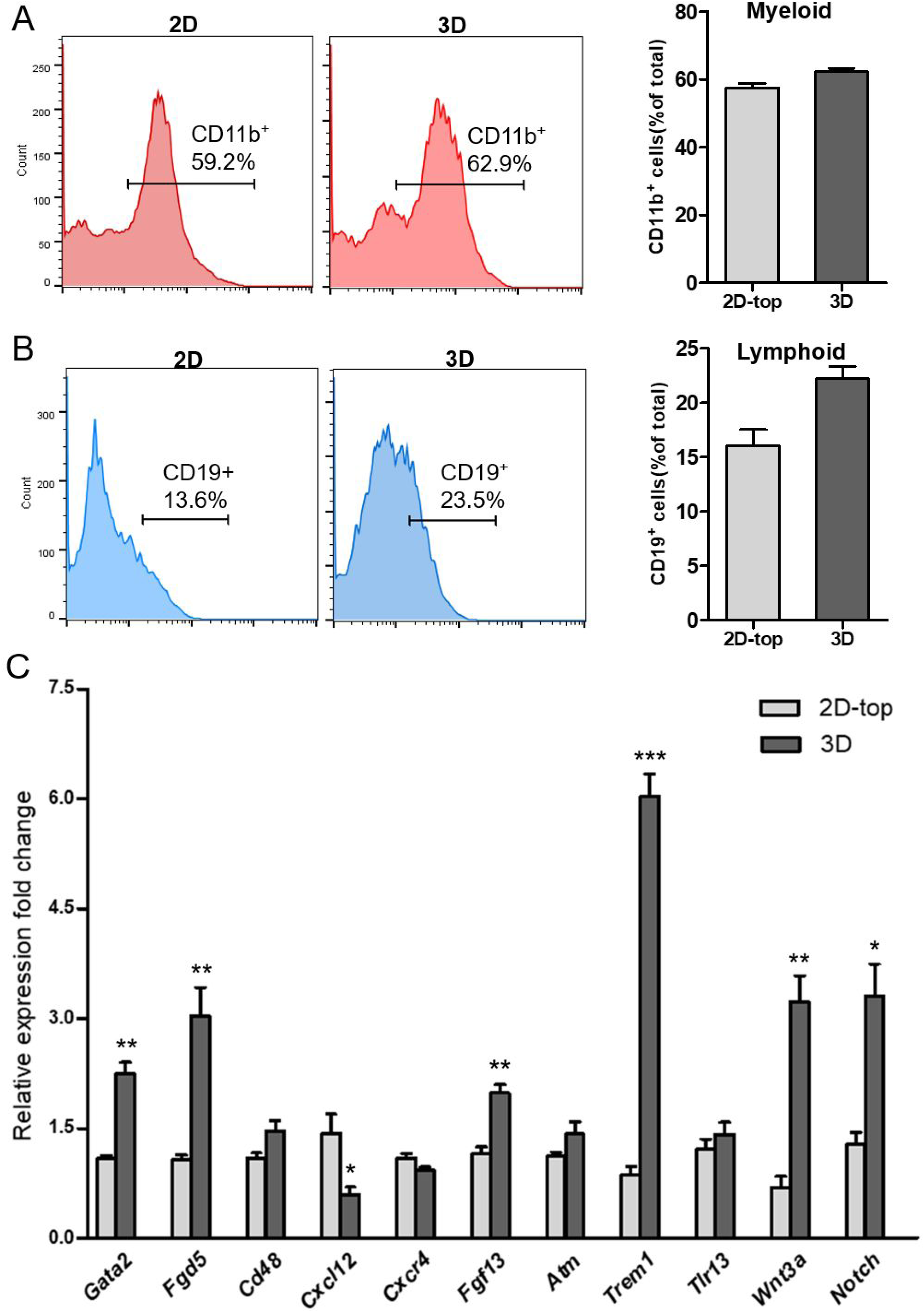
Effect of dimensionality (2D versus 3D) on frequencies of bone marrow myeloid (CD11b^+^) and lymphoid (CD19^+^) cells. Repeated FACS analysis had the same trend. A) Flow cytometry analysis for quantifying percentage of CD11b^+^ cells. The increase showed statistical difference (p=0.0350). (B) A significant elevation in the percentage of CD19^+^ cells cultured in 3D matrix (p=0.0277). n=3. Data are presented as the mean ± SEM. *p<0.05. **(C)** Bar plots graph showing mRNA fold changes through RT-qPCR experiment. Selected genes are differentially expressed in 2D and 3D platforms. *Trem1* gene with the largest increase in 3D was demonstrated. Error bar indicates standard deviation. n = 6. *p<0.05,**p<0.01,***p<0.0001.

ECM substrate elasticity between 1 kPa and 196 kPa has been revealed to affect HSCs viability and cell morphologies(11, 12). 2D hydrogel with 3.7kPa and 44kPa, which respectively mimic tissue rigidity of microenvironment closing to sinusoid(~3kPa) and bone surface(~40 kPa), were demonstrated to direct fate decisions of early HSCs(13). To determine HSCPs have mechanical perception to softer matrix (<100Pa), we used collagen I to fabricate encapsulation hydrogel culture system with defined mechanical property. Collagen is the most abundant protein in mammalian tissues and can self-aggregate to form stable fibers basing on the structure featured by three polypeptide chains wrap around one another to compose a three-stranded rope(14). Excellent biocompatibility and safety, make different types of collagen the primary resource in biomedical applications. Gel stiffness can represent the range of elastic modulus of adipose-rich BM cavity (0~1kPa)(12). Expansion of Lin^−^ hematopoietic progenitors shows stiffness dependence, suggesting when response to matrix stiffness, HSPCs appear mechano-sensitivity, even in a lower elastic modulus range. Dimensionality cues also been mentioned to associate with stem cell fate determination. Previous studies only study changes on cell viability and spread areas induced by system dimensionality(12). Hence, we next asked whether matrix-dimensionality play roles in preserving hematopoietic cell pool. In our study, 3D encapsulation culture method conduce to maintain the cell proliferation ability. Percentage of Lin^−^ hematopoietic progenitors cultured in 3D system is more than 2.5 folds higher than those cultured on matrix surface. Embedded LSK cells number and percentage are also drastically higher. In addition, embedded environment is benefit to the differentiative capacity of hematopoietic cells in vitro. Meanwhile, the function link of external physical signals and internal molecular mechanism are demonstrated. We can speculate that environment dimensionality may be directly sensed by BM HSPCs, and then external signal give rise to the transcriptional activation to mediate cell growth and differentiation. ***Trem1***, which has been characterized as a mature stage of myeloid development in adult hematopoiesis, plays a pivotal role in inflammatory reaction(15, 16). Our result reflects dimensionality of culture environment is particularly connected with myeloid lineage retention. But, degradation of Cxcr4 and Cxcl12 levels is confirmed. Hematopoietic cells in completely encapsulated state probably have impaired migration(17).

We provide evidence that 3D cell-matrix interaction provide necessary information to retain the growth of hematopoietic stem progenitor cells. After all, 2D system can’t stimulate the real physical situation of native tissue in vivo, whereas 3D niche-mimicry system enforce the cell-microenvironment interaction by stabilizing cell-environment mechano-sensing(18). Unlike other 3D platforms those do not support infiltration of the cells inside the material(such as 3D nanofiber platform), collagen encapsulating platform improves the exposure area of cells inside the complex polymeric fiber networks to the external environment, ensuring a true 3D system to promote several cellular mechanisms including cell adhesion, motility, and mechano-sensing(9, 18). We consider dimensionality is a critical design element for artificial niche aimed at resolving some first order questions about the biological processes of HSC. But the present research has its limitations. Because the BM progenitor cells we used included the hematopoietic cells and other BM niche cells, more later experiments need to perform to identified whether the system dimensionality regulated the fate of HSCs independently or in a combination with other niche cells. Of course, further molecular mechanisms reacting to the external physical cues also need to be revealed, basing on our first term study.

## Author Contributions

Hui Yang, Pan Zhang designed research; Yang Hui provided financial support; Pan Zhang performed research and wrote the paper; Chen Zhang and Jiyang Han analyzed data; Jingsong Gao and Wenjuan Zhao participated in paper writing.

## Acknowledgments

This work was supported by grants from National Natural Science Foundation of China (NSFC number 11722220 and 11672246).

